# Deep Feature Extraction for Resting-State Functional MRI by Self-Supervised Learning and Application to Schizophrenia Diagnosis

**DOI:** 10.1101/2020.08.22.260406

**Authors:** Yuki Hashimoto, Yosuke Ogata, Manabu Honda, Yuichi Yamashita

## Abstract

In this study, we propose a novel deep-learning technique for functional MRI analysis. We introduced an “identity feature” by a self-supervised learning schema, in which a neural network is trained solely based on the MRI-scans; furthermore, training does not require any explicit labels. The proposed method demonstrated that each temporal slice of resting state functional MRI contains enough information to identify the subject. The network learned a feature space in which the features were clustered per subject for the test data as well as for the training data; this is unlike the features extracted by conventional methods including region of interests pooling signals and principle component analysis. In addition, using a simple linear classifier for the identity features, we demonstrated that the extracted features could contribute to schizophrenia diagnosis. The classification accuracy of our identity features was higher than that of the conventional functional connectivity. Our results suggested that our proposed training scheme of the neural network captured brain functioning related to the diagnosis of psychiatric disorders as well as the identity of the subject. Our results together highlight the validity of our proposed technique as a design for self-supervised learning.

## 1. Introduction

In this study, we propose a novel deep-learning technique which extracts a feature from brain functional magnetic resonance images (fMRIs). Our proposed method solely depends on MRI-scans and does not require any additional data regarding the subjects (e.g., diseases or cognitive impairments). The extracted features effectively capture the psychopathological characteristics of the subjects. Recent advances in machine learning have demonstrated its capability for medical sciences (Esteva et al., 2017; De Fauw et al., 2018), and skin cancers have been successfully diagnosed from skin images (Esteva et al., 2017) and retinal diseases from three-dimensional optical coherence tomography (OCT) images (De Fauw et al., 2018). In addition, Titano et al. (2018) reported that machine learning with three-dimensional brain computed tomography (CT) images performed well in terms of detection of acute neurologic events including stroke, hemorrhage, and hydrocephalus. These studies suggested the further potential of the deep neural networks, especially for the analysis of spatially structured data, including MRIs and functional MRIs. These studies trained a neural network to directly infer diseases from the input. This framework is called fully supervised learning and is known to be effective when a large training dataset with accurate labels is available. Titano et al. (2018), who aimed to classify acute neurological events, collected 37,236 brain images with clinical annotations for training and used 96,303 extra clinical reports to make the clinical annotations more suitable for training. In the supervised leaning framework, the network is specialized for the target diseases, which further enhances the performance.

However, the requirement of a vast amount of training data is not always practical; the number of patients is sometimes too small to train a neural network (Durstewitz et al., 2019; Khosla et al., 2019), and the accurate diagnoses require expert skills (Durstewitz et al., 2019). These drawbacks are remarkable especially for psychiatric disorders because the sample size tends to be small, accurate diagnoses are especially difficult, and the underlying mechanisms are still under discussion. In contrast, self-supervised learning does not require any explicit labels for training. Instead, the teacher signals (i.e., labels) are generated from the original input data in self-supervised learning. For example, Noroozi and Favoro (2016) proposed a self-supervised learning scheme for natural image processing, in which the input image was divided into nine pieces, and the network was trained to infer the original position of each piece. The intermediate outputs of the network were subsequently fed into another linear classifier, which resulted in comparable performance to fully supervised deep neural networks. The advantages of self-supervised training potentially overcome the shortages of clean labels for psychiatric disorders, although the teacher signal must be carefully designed. This study proposes the design of a self-generated teacher signal for resting-state functional MRI; we used the temporal slices as input, and the subject ID as the teacher signal. The explicit labels enable the network to generate a compact feature that represents a conceptual distance from the owner of the input to the subjects used in the training. In this study, we experimentally showed that (i) each temporal slice of functional MRI contains enough information to identify the subject, (ii) the network learned a feature space in which the features cluster subject-by-subject for test data as well as for training data, and (iii) the extracted feature contributes to schizophrenia diagnosis. These experiments together exhibit the validity of our proposed method as design for self-supervised learning.

## 2. Materials and Methods

### 2.1. Dataset

We used a dataset from The Center for Biomedical Research Excellence (COBRE) (The Mind Research Network & The University of New Mexico, 2012). The dataset is composed of anatomical and resting-state functional MRI scans; 72 scans were from schizophrenia patients and 75 from healthy controls. The anatomical and functional scans were acquired by MPRAGE and EPI. Each functional scan was composed of 150 timepoints, and the repetition time was two seconds. Each timepoint was originally composed of 64 × 64 × 32 voxels, whose size was 3 × 3 × 4 mm^3^. We excluded subjects without meta-data and controls with other psychiatric diseases, resulting in 69 patients and 72 controls. We randomly divided the patients and controls into training 1, training 2, and test dataset. The training 1 dataset was used for training the neural network, and training 2 was used for training the linear regressor for inferring the subject attributes. The number of patients *p* and controls *c* was (*p*, *c*) = (51, 54) in training 1, (9, 9) in training 2, and (9, 9) in test datasets. These numbers were determined in advance, and then, the subjects were randomly allocated to each group.

### 2.2. Preprocessing

Each scan was spatially realigned, timing-corrected, segmented, normalized to MNI coordinate, and smoothed by the default pre-processing pipeline of the CONN toolbox (Whitfield-Gabrieli and Nieto-Castanon, 2012). The resulting spatial resolution was (91, 109, 91). Subsequently, its default denoising pipeline was applied. We masked out voxels of white matter and cerebrospinal tissue and acquired a gray matter mask of 150,571 voxels. We then normalized the signal intensity within the gray matter mask for each scan to prevent the network from learning the bias and variance as a clue for subject classification. The signals out of the gray-matter mask were replaced by zero.

### 2.3. Training 1

The input of the network was a batch of temporal MRI slices, whose size was set to (80, 96, 80) by trimming the zero-filled region. The network included four convolutional blocks, followed by two convolutional layers and one dense layer. Each block consisted of two three-dimensional convolutions and one average pooling layer (Fig. 1).

**Figure 1.**
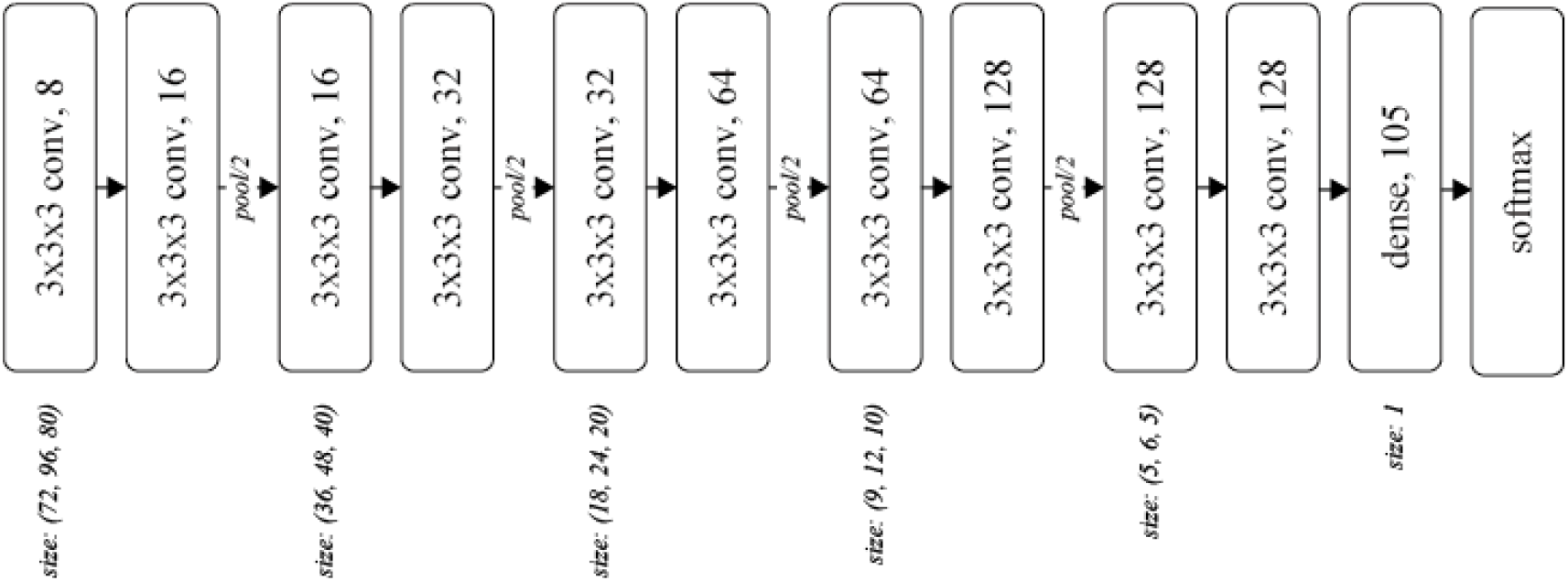
Network architecture. Each rounded square represents a layer with the weight parameters. The number after the comma denotes the number of channels for the layer.

The kernel size *k* and stride *s* were (*k*, *s*) = (3, 1) for each convolutional layer, and (2, 2) for each pooling layer. The number of output channels was set as 8 at the first convolutional layer and doubled before the pooling layers, resulting in 128 before the dense layer. We used softmax cross-entropy as the loss function, which was computed against a 105-dimensinal one-hot vector of subject ID. The network was optimized using Adam (Kingma and Ba, 2015) with *α* =00.0001 for the first 17,000 iterations and *α*=00.00001 for the following 110,000 iterations, with a batch size of 32.

### 2.4. Training 2

The output of the dense layer was extracted for each timepoint as a feature vector. Subsequently, the feature vectors were averaged for each subject, yielding an *identity feature* for each subject. The identity features for the training 2 dataset were then fed into train linear classifier (regressor) for learning schizophrenia history and age.

We also trained a linear classifier with a slightly modified version of the feature vector, in which the average of all elements in the feature vector was subtracted from each element. This operation was naturally introduced by the formulation of the softmax function, in which the subtraction of the average does not affect the output of the function or the training process. In the following sections, we call the original feature vector (the output of the dense layer) as *classification*, and the modified one as *classification+*.

### 2.5. Experiment 1: training convergence

The training accuracy of the subject classification was computed to evaluate training. Reporting training accuracy is slightly unconventional in studies on neural networks because the convergence of training is now trivial in conventional two-dimensional natural scene image processing. However, to the best of our knowledge, this is the first report which trained networks to classify the subject from a single timepoint of functional MRI by stacked three-dimensional convolutions, and we concluded that the training convergence is worthy enough to report.

### 2.6. Experiment 2: qualitative analysis of extracted features

The characteristics of the acquired feature space were first qualitatively analyzed. We plotted the feature vectors in the training 2 and test datasets by t-distributed stochastic neighbor embedding (t-SNE) (Maaten and Hinton, 2008). The clusters were then quantitatively evaluated by precision@150 for each identity feature. Because the number of timepoints was 150 for each subject, precision@150 would be 1 if all the feature vectors for a subject clustered around his identity feature. The formula of precision@150 is given in the appendix. We applied these qualitative and quantitative analyses to the features of the *classification* and *classification+* feature vectors as well as the signals averaged over the region of interest (ROI) defined by the automated anatomical labeling (AAL) atlas (Suk et al., 2016; see Tzourio-Mazoyer et al., 2002 as the reference to AAL), and the top three and 10,000 principle components, which are the bases of independent components (Damaraju et al., 2014; Beckmann et al. (2004) for the relation of the principle and independent components in the linear space).

### 2.7. Experiment 3: relation to subject’s attributes

The schizophrenia classifier and age regressor developed in Section 2.4 were applied to the test dataset. The classification accuracy was computed and tested using a sign test. This procedure was also applied to the identity features of the ROI-pooled signals, the top three and 10,000 principle components, similar to that in Experiment 2. In addition, the procedure was applied to the functional connectivity matrix (Liang et al., 2006; Kim et al., 2016), defined as the correlation coefficients among the time series of the ROI-pooled signals.

### 2.8. Ethics statement

All experiments in this study were performed in accordance with the Ethical Guidelines for Medical and Health Research Involving Human Subjects in Japan.

## 3. Result

### 3.1. Experiment 1: training convergence

The network was trained to classify the subject ID from each time point of fMRI. The training accuracy at the 127,000 iteration was 97.85%, which was considerably improved over the chance rate, suggesting that the training successfully converged.

### 3.2. Experiment 2: qualitative analysis of extracted features

The distributions of the feature vectors extracted by our proposed neural network, ROI-pooling, and PCA were visualized by t-SNE, and are depicted in Fig. 2.

**Figure 2.**
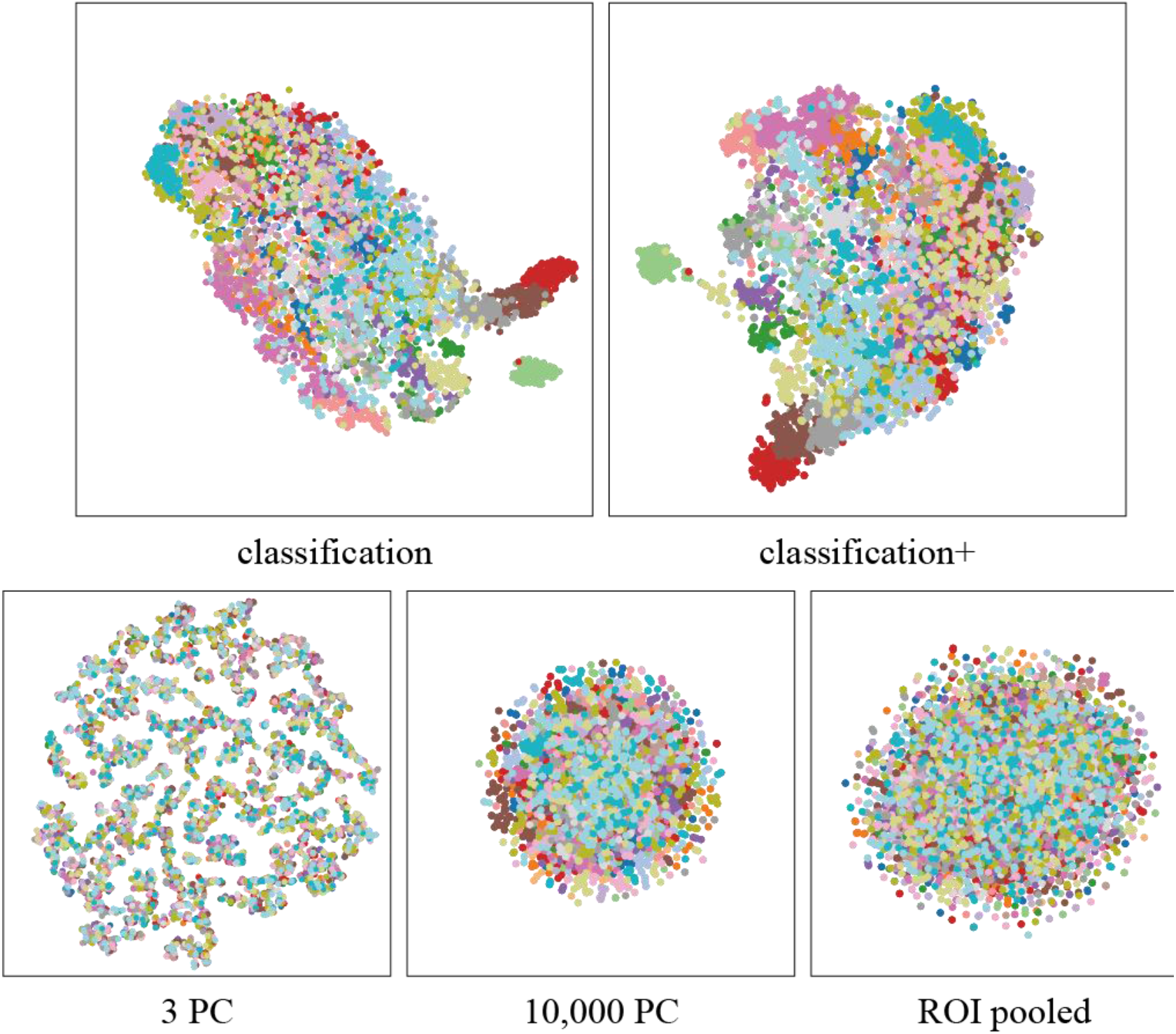
Distribution of feature vectors, visualized by t-SNE. Each dot represents a feature vector for a single timepoint, colored for each subject.

The features extracted by the network clustered for each subject, unlike the features extracted by ROI-pooling and PCA. The clustering performance was quantitatively evaluated using precision@150 around the identity feature for each subject. The precision@150 was 81.5% and 61.4% for our proposed *classification* and *classification+* feature vectors, whereas it was 0.56% for the ROI-pooled feature and the top three and 10,000 principle components (Fig. 3).

**Figure 3.**
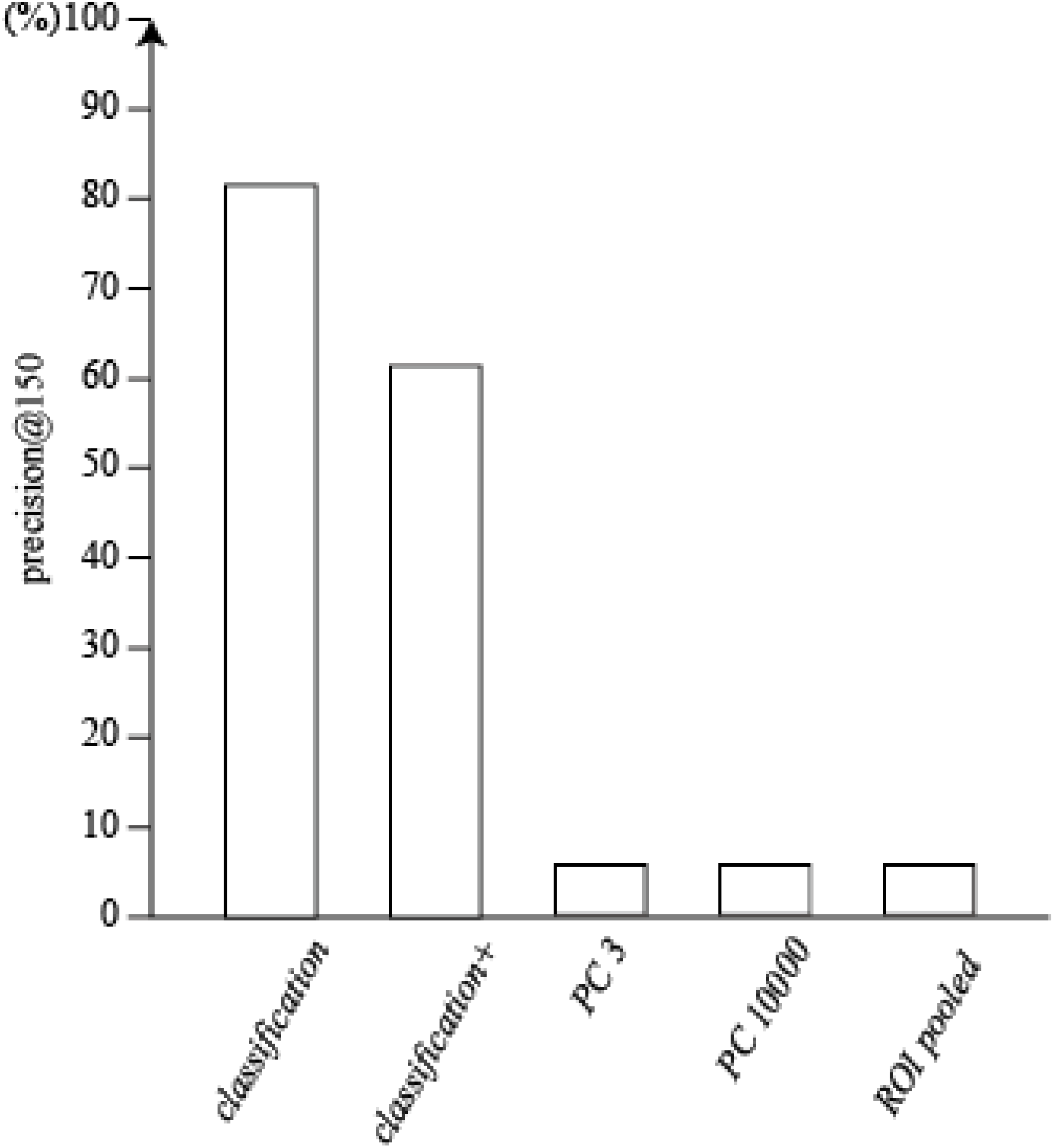
Quantitative index of the cluster for each feature vector. The y-axis represents the precision@150 for the subject clustering task.

### 3.3. Experiment 3: schizophrenia diagnosis

The average of the features was computed as the identity feature for each subject, and the identity features were fed into a linear classifier with a logistic loss function. The accuracies were 61.1 and 77.8% for the identity feature of our proposed *classification* and *classification+* feature vectors, respectively. The performance of *classification+* was significantly better than the chance (p 0 0.015). The accuracy was 72.2% for the connectivity matrix, which was marginally higher above the chance (p 0 0.048). The identity features of the top three and 10,000 principle components and the ROI-pooled signals did not significantly discriminate the schizophrenia and control group (acc. 0 27.8%, 50%, and 61.1%, respectively), as shown in Fig. 4.

**Figure 4.**
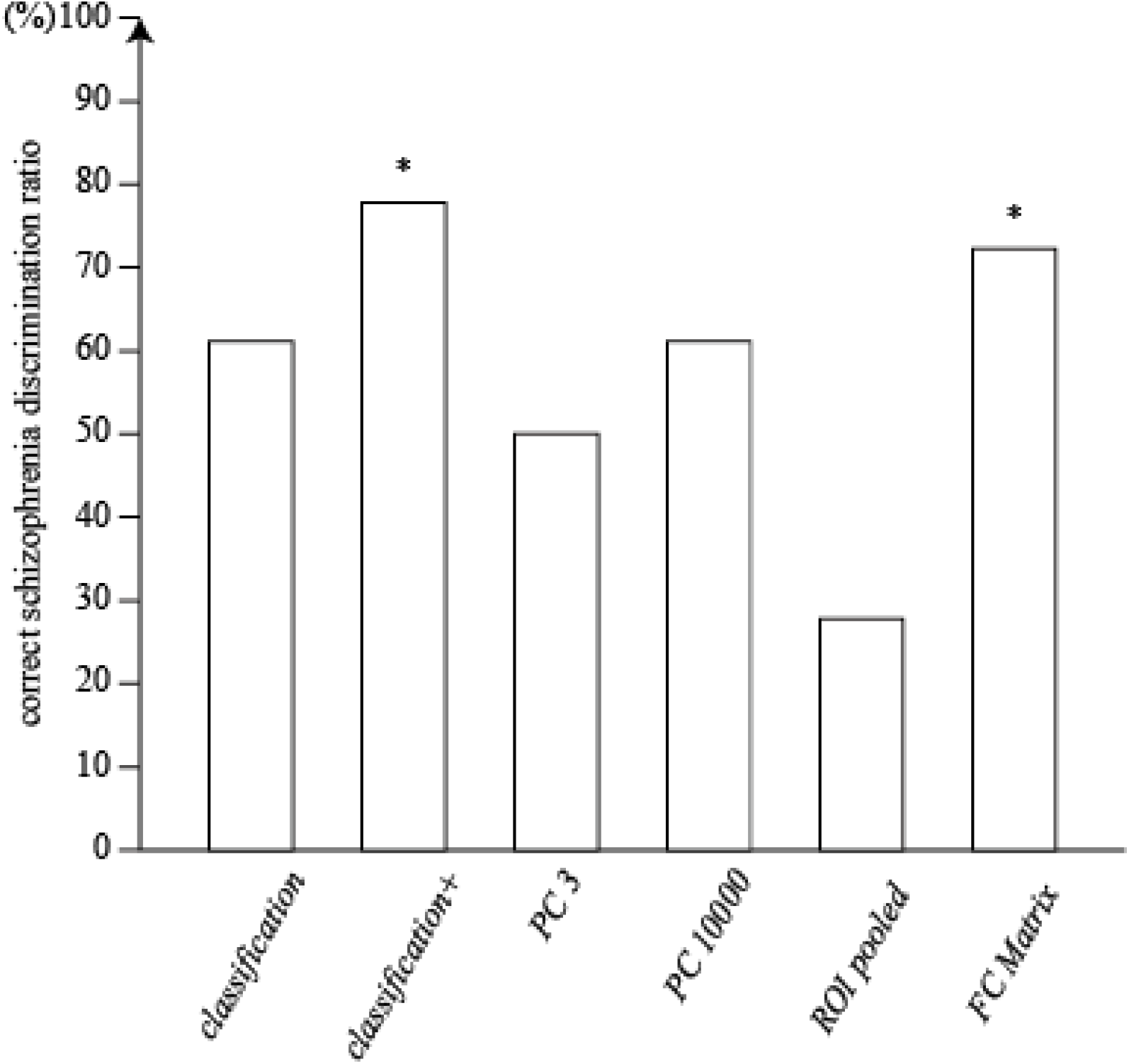
Accuracy of the schizophrenia discrimination. The single asterisks show the statistical significance at *α*=00.05.

Similarly, subject age was regressed from the identity feature. The correlations between the predicted and actual age were not significant (r=0.128 and 0.115 for *classification* and *classification+)*, while the top three principle components showed significant correlation (r=0.57, p=0.013). The other conditions (i.e., the top 10,000 principle components, ROI-pooled signals, and functional connectivity matrix) did not showed significant correlation (r=−0.21, −0.29, and 0.34, respectively), as shown in Fig. 5.

**Figure 5.**
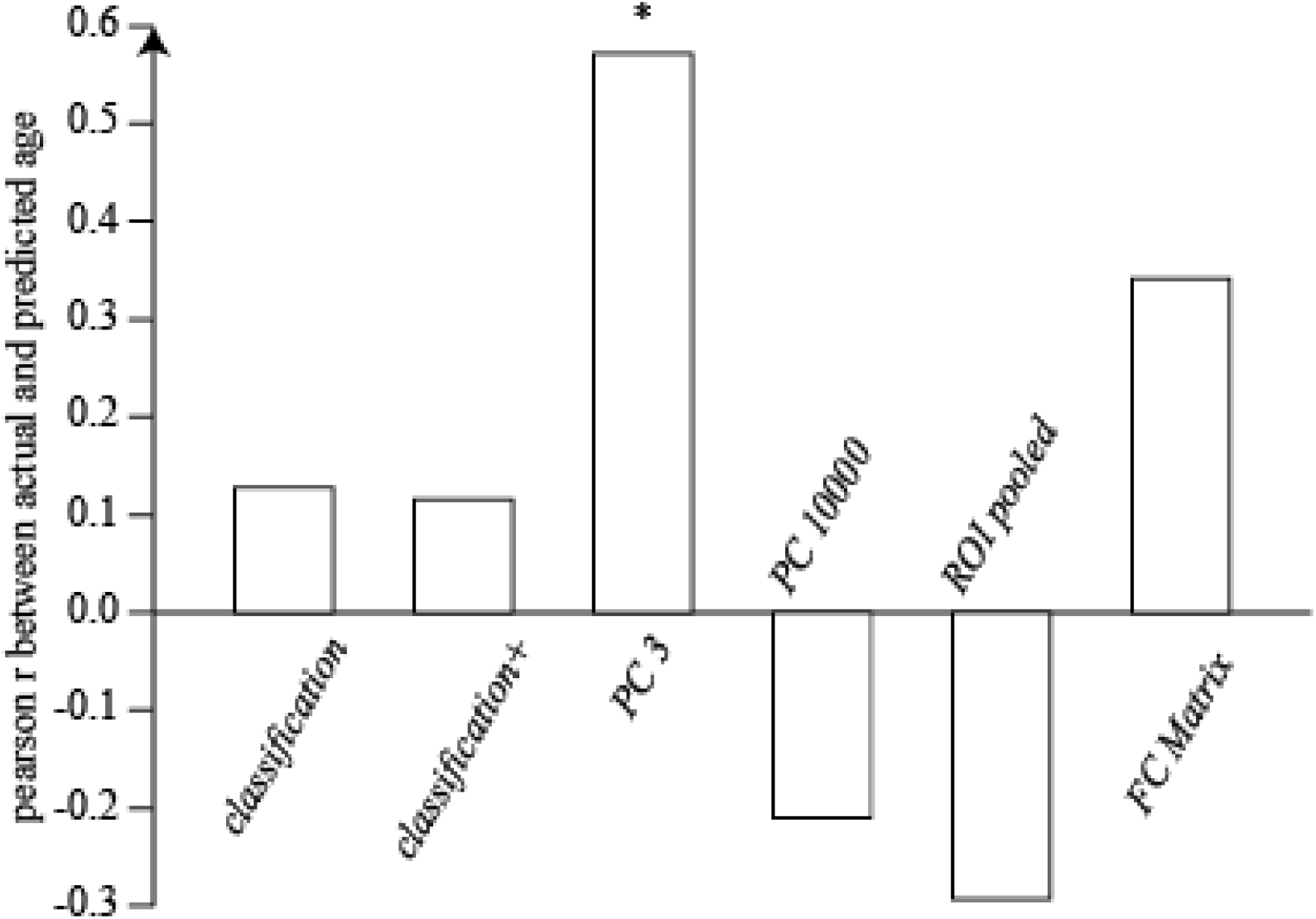
Correlation between the actual and predicted age for each feature vector. The asterisk indicates a significant correlation at of *α*=00.05.

## 4. Discussion

We have shown that (i) the self-supervised learning scheme let our neural network to acquire the projection from the high (~10^6^) dimensional signal space to the lower dimensional (~10^2^) feature space in which each dimension represented subject identity in the training dataset, (ii) the capability of the subject identification was generalized to the unknown subjects in the test dataset, and (iii) the temporal average of the extracted feature vector reflected the psychiatric status of the subjects. Surprisingly, our proposed method performed even better than the functional connectivity matrix for schizophrenia diagnosis, which has been regarded as a promising biomarker of cognitive functions (Liang et al., 2006; Kim et al., 2016).

The transferred capability from the subject identification to schizophrenia diagnosis can be regarded as a kind of “deep feature extraction”. In the natural scene image processing, the intermediate output in a neural network pre-trained with a large-dataset classification often works well in another task, known as a “deep feature extraction” (Oquab et al., 2014). The underlying mechanism of the transferability is still under debate; however, one of the dominant hypotheses is that the stacked two-dimensional convolution itself works as the statistical prior of the natural scene images, regardless of the training task (Ulyanov et al., 2018). Our results showed that the transference also occurred with the combination of the human-brain T2* images and the stacked three-dimensional convolutions.

Our feature did not correlate with the subject’s age, unlike the psychiatric status. This result suggests that subjects with similar psychiatric status are adjacent on the feature space, whereas similar age subjects are not. Given this discussion, the linear-decomposition-based features (i.e. the principle / independent components) and the functional connectivity matrix might have potentially ignored the discontinuity on the signal-space, yielding the results in the subject’s age regression different from our identity feature.

The superiority of our feature over the functional connectivity in schizophrenia discrimination might be attributed to the local interactions of the signal. In the functional connectivity analysis, the signals are averaged for each ROI, discarding the local signal interactions. In contrast, previous studies have reported that both global and local activities in brain lead to our cognitive functions (see Panzeri, Macke, Gross & Kayser, 2015 for review). Both of the local and global interactions are modeled in the neural network, and it might have led to a positive effect in schizophrenia discrimination.

In this study, we introduced a novel self-supervised learning scheme and highlighted some of the characteristics of the extracted feature, especially in terms of the relation to schizophrenia. However, there are many questions not answered in this study. For example, the optimal number of subjects in the training, the optimal neural network architecture, the more detailed relations between the feature and the subject’s attributes, and the mathematical analyses about the feature space. We hope these will be uncovered in future works along the further accumulation of available datasets and with the advancement in the field of machine learning.

## Appendix Formula for precision@K

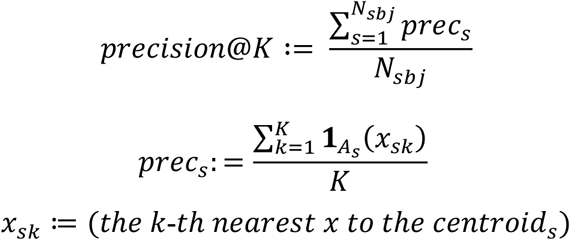

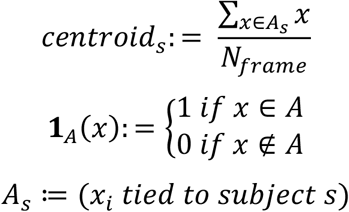

## Acknowledgments

This work was partially supported by JSPS KAKENHI [grant numbers JP17H06039, JP18KT0021, JP18K07597, JP19H04998, and JP20H00625] and JST CREST [grant number JPMJCR16E2].

## Data availability

The primary data can be obtained from public databases, the Centers for Biomedical Research Excellence (COBRE; http://fcon_1000.projects.nitrc.org/indi/retro/cobre.html).

## Author contributions

YH and YY conceived and designed the research. YH conducted the experiments and analyzed the data. YO supported preprocessing of MRI data. YH and YY drafted the manuscript. YO and MH provided critical revisions. All authors contributed to and have approved the final manuscript.

## References

Damaraju, E., Allen, E.A., Belger, A., Ford, J.M., McEwen, S., Mathalon, D. H., Mueller, B.A., Pearlson, G.D., Potkin, S.G., Preda, A., Turner, J.A., Vaidya, J.G., van Erp, T.G., Calhoun, V.D., 2014. Dynamic functional connectivity analysis reveals transient states of dysconnectivity in schizophrenia. NeuroImage: Clinical, 5, 298–308.

De Fauw, J., Ledsam, J. R., Romera-Paredes, B., Nikolov, S., Tomasev, N., Blackwell, S., Askham, H., Glorot, X., O’Donoghue, B., Visentin, D. and van den Driessche, G., 2018. Clinically applicable deep learning for diagnosis and referral in retinal disease. Nat. Med. 24 (9), 1342–1350.

Durstewitz, D., Koppe, G., Meyer-Lindenberg, A., 2019. Deep neural networks in psychiatry. Mol. Psychiatry 24 (11), 1583–1598.

Estava, A., Kuprel, B., Novoa, R.A., Ko, J., Swetter, S. M., Blau, H.M., Thrun, S., 2017. Dermatologist-level classification of skin cancer with deep neural networks. Nature, 542 (7639), 115–118.

Khosla, M., Jamison, K., Ngo, G.H., Kuceyeski, A., Sabuncu, M.R., 2019. Machine learning in resting-state fMRI analysis. Magn. Reson. Imaging, 64, 101–121.

Kim, J., Calhoun, V.D., Shim, E., Lee, J.H., 2016. Deep neural network with weight sparsity control and pre-training extracts hierarchical features and enhances classification performance: Evidence from whole-brain resting-state functional connectivity patterns of schizophrenia. Neuroimage 124 (A), 127–146.

Kingma, D.P., Ba, L.J., 2015. Adam: A method for stochastic optimization [Conference presentation abstract]. International Conference on Learning Representations, San Diego, CA, United States. https://arxiv.org/abs/1412.6980v5

Liang, M., Zhou, Y., Jiang, T., Liu, Z., Tian, L., Liu, H., Hao, Y., 2006. Widespread functional disconnectivity in schizophrenia with resting-state functional magnetic resonance imaging. NeuroReport 17 (2), 209–213.

Maaten, L. V. D., Hinton, G., 2008. Visualizing data using t-SNE. J. Mach. Learn. Res. 9, 2579–2605.

Noroozi, M., Favaro, P., 2016. Unsupervised learning on visual representations by solving jigsaw puzzles, In Lecture notes in computer science, in: Leibe, B., Matas, J., Sebe, N. Welling, M. (Eds.). Cham: Springer, 69–84.

Oquab, M., Bottou, L., Laptev, I., Sivic, J., 2014. Learning and transferring mid-level image representations using convolutional neural networks. Proceedings of the IEEE conference on computer vision and pattern recognition. IEEE 1717–1724.

Panzeri, S., Macke, J. H., Gross, J., Kayser, C., 2015. Neural population coding: Combining insights from microscopic and mass signals. Trends Cogn. Sci. 19 (3), 162–172.

Suk, H. I., Wee, C. Y., Lee, S. W., Shen, D., 2016. State-space model with deep learning for functional dynamics estimation in resting-state fMRI. Neuroimage 129, 292–307.

The Mind Research Network & the University of New Mexico. (2012) The Center for Biomedical Research Excellence. http://fcon_1000.projects.nitrc.org/indi/retro/cobre.html.

Titano, J. J., Badgeley, M., Schefflein, J., Pain, M., Su, A., Cai, M., Swinburne, N., Zech, J., Kim, J., Bederson, J., Mocco, J., Drayer, B., Lehar, J., Cho, S., Costa, A., Oermann, E. K., 2018. Automated deep-neural-network surveillance of cranial images for acute neurologic events. Nat. Med. 24 (9), 1337–1341.

Tzourio-Mazoyer, N., Landeau, B., Papathanassiou, D., Crivello, F., Etard, O., Delcroix, N., Mazoyer, B., Joliot, M., 2002. Automated anatomical labeling of activations in SPM using a macroscopic anatomical parcellation of the MNI MRI single-subject brain. Neuroimage 15 (1), 273–289.

Ulyanov, D., Vedaldi, A., & Ulyanov, D., 2018. Deep image prior. Proceedings of the IEEE conference on computer vision and pattern recognition. IEEE 9446–9454.

Whitfield-Gabrieli, S., Nieto-Castanon, A., 2012. Conn: a functional connectivity toolbox for correlated and anticorrelated brain networks. Brain Connect. 2 (3), 125–141

